# SARS-CoV-2 variants show temperature-dependent enhanced polymerase activity in the upper respiratory tract and high transmissibility

**DOI:** 10.1101/2022.09.27.509689

**Authors:** Se-Mi Kim, Eun-Ha Kim, Mark Anthony B. Casel, Young-Il Kim, Rong Sun, Mi-Jeong Kwack, Ji-Seung Yoo, Min-Ah Yu, Kwang-Min Yu, Seung-Gyu Jang, Rare Rollon, Jeong Ho Choi, JuRyeon Gil, Kiyoung Eun, Hyunggee Kim, Armin Ensser, Jungwon Hwang, Min-Suk Song, Myung Hee Kim, Jae U Jung, Young Ki Choi

## Abstract

With the convergent global emergence of SARS-CoV-2 variants of concern (VOC), a precise comparison study of viral fitness and transmission characteristics is necessary for the prediction of dominant VOCs and the development of suitable countermeasures. While airway temperature plays important roles in the fitness and transmissibility of respiratory tract viruses, it has not been well studied with SARS-CoV-2. Here we demonstrate that natural temperature differences between the upper (33°C) and lower (37°C) respiratory tract have profound effects on SARS-CoV-2 replication and transmission. Specifically, SARS-COV-2 variants containing the P323L or P323L/G671S mutation in the NSP12 RNA-dependent RNA polymerase (RdRp) exhibited enhanced RdRp enzymatic activity at 33°C compared to 37°C and high transmissibility in ferrets. MicroScale Thermophoresis demonstrated that the NSP12 P323L or P323L/G671S mutation stabilized the NSP12-NSP7-NSP8 complex interaction. Furthermore, reverse genetics-derived SARS-CoV-2 variants containing the NSP12 P323L or P323L/G671S mutation displayed enhanced replication at 33°C, and high transmission in ferrets. This suggests that the evolutionarily forced NSP12 P323L and P323L/G671S mutations of recent SARS-CoV-2 VOC strains are associated with increases of the RdRp complex stability and enzymatic activity, promoting the high transmissibility.

## Introduction

The rapid evolutionary rate among RNA viruses is associated with their high mutation rates. SARS-CoV-2 exhibited evolutionary stasis at the beginning of the COVID-19 pandemic; however, the virus rapidly acquired mutations despite the proofreading ability of its RNA-dependent RNA polymerase (RdRp) ^1,2^. Although most mutations are reportedly deleterious or neutral towards a virus ^3^, the presence of selective pressure has contributed to the sporadic emergence of genetically diverse SARS-CoV-2 variants. Accordingly, at least six different SARS-CoV-2 clades have been reported ^4^, followed by the emergence of notable heavily mutated SARS-CoV-2 variants, suggesting the rapid accumulation of genetic mutations during human infections. The D614G mutation in the SARS-CoV-2 spike (S) protein was first reported in early March 2020, and most recent variants have maintained this mutation along with the accumulation of additional mutations or deletions in their genome. The D614G mutated S protein shows high affinity for the host receptor, which relates to the virus’ infectivity and transmission ability ^5–7^. Beside the S protein mutations, P323L is a key mutation in the RdRp encoded by NSP12, which is of particular interest because P323 is within the domain that interacts with the Nsp7-Nsp8 cofactors that regulate viral polymerase function ^8^. Recent studies report that the P323L mutation in NSP12 is associated with the mutation rate in the viral genome ^9^ and with selection pressure advantage compared to the P323 virus ^10^. However, only limited detailed functional studies have examined the P323L-NSP12 mutation. Furthermore, among multiple SARS-CoV-2 variants, B.1.1.529 (Omicron) was most recently declared a new variant of concern (VOC) by the WHO ^11^, and Omicron and its sub-variants (including BA.2.75) are co-circulating globally and showing dynamic transmission patterns. Genomic surveillance and epidemiological investigations have demonstrated that mainly the S protein mutations in these recent variants may contribute to their rapid replication and increased transmissibility in humans ^12–17^; however, there has not yet been a systematic comparison of the full importance of the mutations among variants.

During multiple transmission cycles in a large population, many RNA viruses may undergo evolutionarily forced genomic mutations that affect viral fitness and transmissibility. With the convergent global emergence of multiple SARS-CoV-2 variants, a precise comparison study of viral fitness and transmission characteristics is necessary to determine the various mutations causing the different biological characteristics of each SARS-CoV-2 variant, to devise a strategy for the continuing development of suitable countermeasures, and to predict which variant may rise to dominance. In the current study, to examine the phenotypic characteristics of each reference strain, we utilized systematic in vitro experimental approaches to assess the biological effects of amino acid mutations in the RdRp. We further used an in vivo ferret model for comparisons of virus replication and transmission kinetics.

## Results

### In vitro growth kinetics and in vivo transmission characteristics of SARS-CoV-2 variants

Epidemiological investigations show that recent variants exhibit increased replication and enhanced transmissibility in humans ^15–17^. To compare growth properties and transmission dynamics among SARS-CoV-2 variants, we selected representative strains based on full-length sequence information from clinical SARS-CoV-2 isolates deposited at the Zoonotic Research Center of Chungbuk National University. Based on the GISAID nomenclature system, the virus isolates were classified as Wuhan-Hu-1-like (L clade, reference strain), P323/D614G (early G clade), Alpha (B.1.1.7), Beta (B.1.351), and Delta (B.1.617.2) variants (table S1). Interestingly, recent SARS-CoV-2 variants harbored two highly abundant mutations, P323L and G671S in the Nsp12 RdRp, with a frequency of 99.7%. The G671S mutation was particularly found among Delta variants. We examined the multi-step growth kinetics of each representative reference virus in Vero, Vero^ACE2/TMPRSS2^, Huh-7, and Calu-3 cells, and found that the Alpha, Beta, and Delta viruses replicated at higher viral titers compared to the reference virus (Fig. 1a-d).

**Fig. 1.**
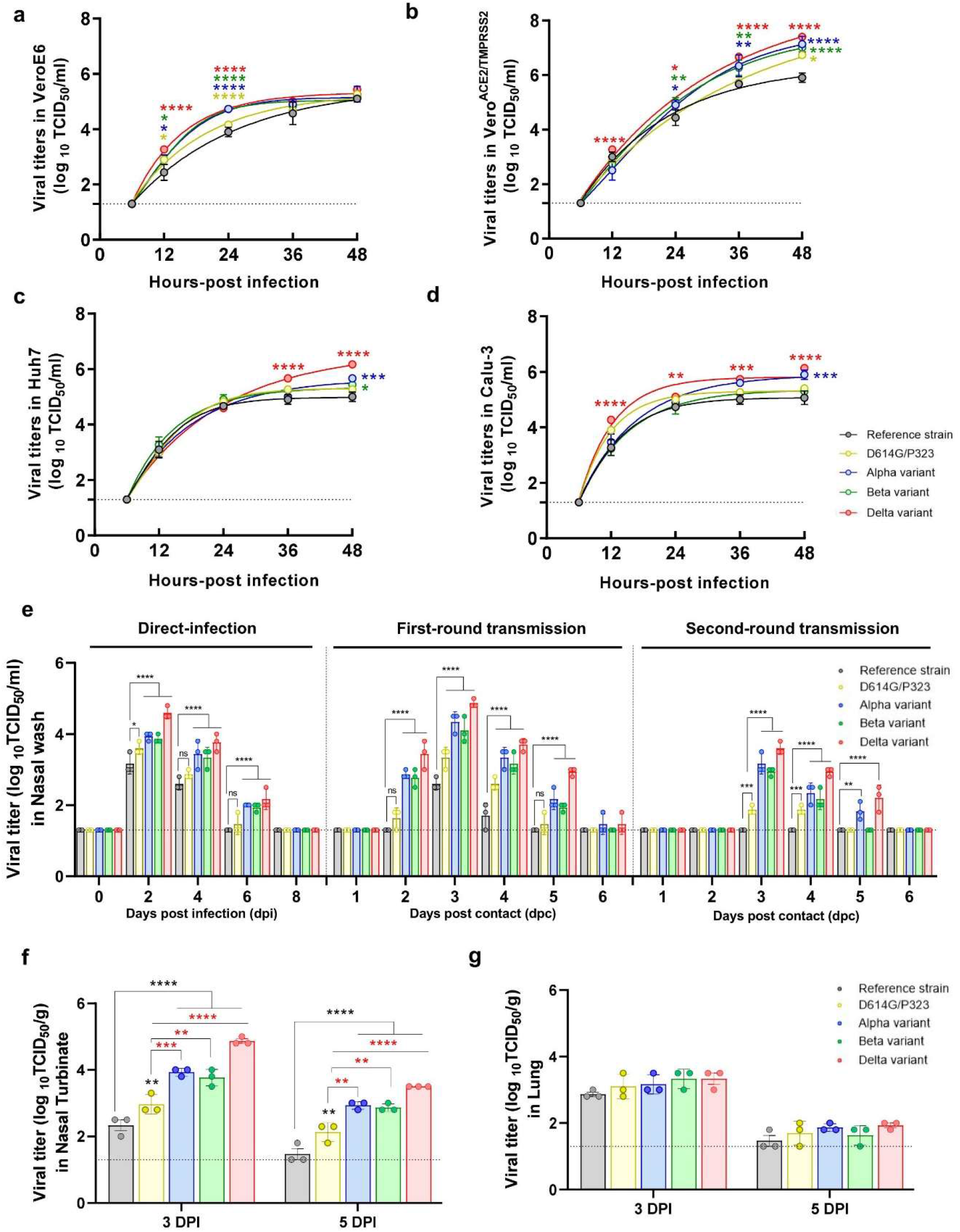
In vitro growth kinetics and in vivo transmission characteristics of SARS-CoV-2 strains and variants in ferrets. Comparison of differential growth kinetics among the reference strain (Wuhan-Hu-1-like, Clade L), D614G strain, and recent variant viruses (Alpha, Beta, and Delta) in **a** Vero, **b** Vero^ACE2/TMPRSS2^, **c** Huh-7, and **d** Calu-3 cells. The cells were infected with viruses at an MOI of 0.01. Ferrets (*n* = 9/group) were inoculated intranasally with 5.0 log_10_ TCID_50_/mL of each representative SARS-CoV-2 strain. Nasal wash specimens were collected at a 2-day interval from directly infected ferrets, and daily from the first-round transmission and second-round transmission ferrets. Ferrets (*n* = 3/group) were sacrificed at 3 and 5 dpi. Infectious virus titers in the **e** nasal wash specimen, **f** nasal turbinates, and **g** lung tissue samples were measured by determining the TCID_50_ per unit of measurement (g or mL) using VeroE6 cells. In **Fig. 1f**, black * indicate statistical results compared to the reference strain, and red **\*** indicate statistical values versus D614G/P323. The limit of detection (1.3 log_10_ TCID_50_/ml or log_10_TCID_50_/g) is indicated by a dotted line for each representation. The *p* values were calculated using a two-way ANOVA, with Turkey’s post-test. \**p* < 0.05; \*\**p* < 0.01; \*\*\**p* < 0.001; \*\*\*\**p* < 0.0001.

To investigate the in vivo differential transmission kinetics of SARS-CoV-2 variants, we conducted a systematic natural transmission study with each representative virus in a ferret infection model, as previously described ^18^. For the initial infection, ferrets (n = 9/group) were infected with 5.0 log_10_ TCID_50_/mL of each representative SARS-CoV-2 strain (Extended Data Fig. 1). The P323/D614G virus showed higher virus titers compared with the reference strain, only at 2 dpi. On the other hand, the Alpha, Beta, and Delta variants showed significantly higher viral titers than the reference strain, from 2 dpi to 6 dpi. Notably, the Delta variant consistently exhibited the highest virus titers from 2 dpi to 6 dpi (Fig. 1e, left panel). For the first-round transmission study, naïve direct contact ferrets (n = 3/group, DC) were co-housed with infected ferrets one day after the initial SARS-CoV-2 infection (Extended Data Fig. 1). From the nasal wash specimens of direct contact ferrets, we recovered infectious P323/D614G, Alpha, Beta, and Delta variants as early as 2 days post-contact (dpc). The Alpha, Beta, and Delta variants consistently exhibited higher virus titers than the reference virus from 2 to 5 dpc (Fig. 1e, middle panel). For the second-round transmission study, three additional naïve ferrets were co-housed with the first-round DC ferrets at 3 dpc (Extended Data Fig. 1). Strikingly, from the nasal washes of the second-round DC ferrets, we detected P323/D614G, Alpha, Beta, and Delta viruses, with a peak viral titer at 3 dpc, while the reference virus was not detected in the second-round DC ferrets at any time-point (Fig. 1e, right panel). Analysis of nasal washes also revealed that P323/D614G virus showed attenuated virus titers compared with other variants, and early clearance by 5 dpc. In both the first- and second-round transmission, the Alpha and Delta variants showed the longest shedding period, until 5 dpc.

To assess the in vivo replication properties of each virus clade in respiratory tissues, we collected nasal turbinates and lungs tissues from directly infected ferrets (n = 3/group) at 3 and 5 dpi. P323/D614G, Alpha, Beta, and Delta variants showed significantly higher virus titers in the nasal turbinates than reference strain, and P323/D614G virus exhibited relatively lower titers compared to the Alpha, Beta, and Delta variants at 3 and 5 dpi (Fig. 1f). In contrast, we observed little or no difference among virus titers in the lung tissues at both time-points (Fig. 1g). Together, these results showed that recent SARS-CoV-2 variants exhibited enhanced growth properties in the upper respiratory tract, as well as rapid and long-term transmission kinetics.

### NSP12 P323L mutation alters SARS-CoV-2 RdRp activity

Besides S protein mutations, viral fitness also considerably affects virus replication, infectivity, and transmissibility among VOCs. Interestingly, the NSP12 P323L mutation has been consistently maintained among SARS-CoV-2 VOCs ^19^. Furthermore, ~99.7% of Delta variants harbor an additional G671S mutation of NSP12 (Extended Data Fig. 2). Since the P323 mutation resides within the NSP12-NSP8 interface ^20^, we performed a cell-based SARS-CoV-2 RdRp activity reporter gene assay, as previously described ^21^. To test how temperature influenced SARS-CoV-2 RdRp activity, the RdRp activity reporter gene assay was conducted at 33°C and 37°C, simulating the upper and lower respiratory tract temperatures, respectively. Although RdRp activity was detectable in the absence of NSP7 and NSP8, the overall RdRp activity was higher in the presence of NSP7 and NSP8 (Fig. 2). Strikingly, the P323L or P323L/G671S NSP12 mutant showed markedly increased RdRp activity at 33°C, compared to wild-type (WT), and this increase was particularly pronounced in 1:2:1 or 2:2:1 ratio condition of the NSP7-NSP8-NSP12 complex (Fig. 2c and d). In contrast, at 37°C, RdRp activity did not significantly differ for the P323L and P323L/G671S NSP12 mutants compared to WT and G671S mutant (Fig. 2). These results indicate that the P323L/G671S mutations have a temperature-dependent synergistic effect on RdRp activity.

**Fig. 2.**
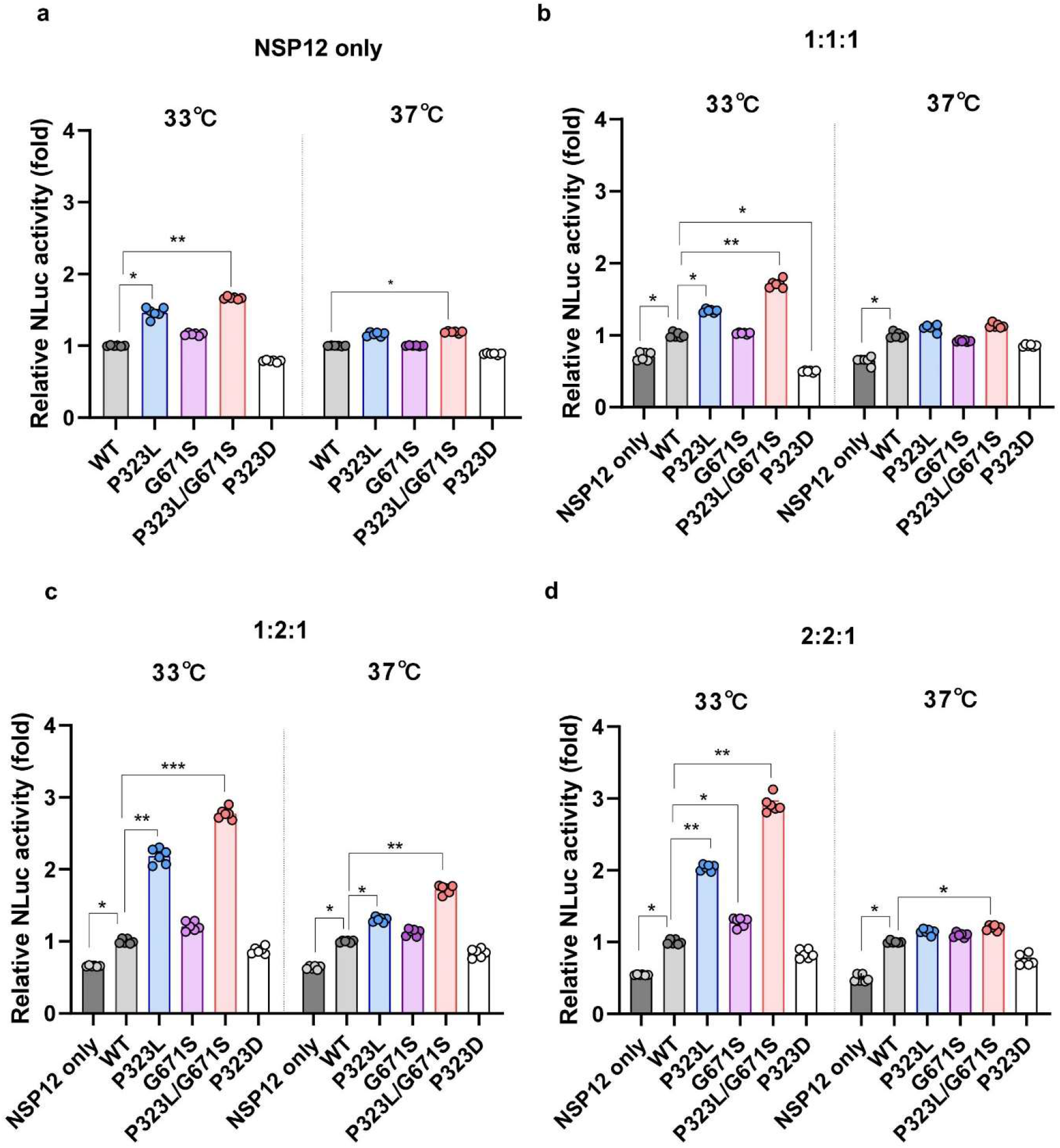
SARS-CoV-2 RdRp reporter gene assay. 293T cells were transiently transfected with SARS-CoV-2 NSP12 wild-type (WT), P323L, G671S, P323L/G671S, and P323D mutant. NSP12 was co-transfected with the NSP7 and NSP8 cofactors, followed by incubation at 33°C or 37°C. **a** NSP12 only. **b** NSP7:NSP8:NSP12 ratio of 1:1:1. **c** NSP7:NSP8:NSP12 ratio of 1:2:1. **d** NSP7:NSP8:NSP12 ratio of 2:2:1. NLuc activities were measured after 24 hours post-transfection. The p values were calculated using a one-way ANOVA, with Dunnett’s post-test. ****p < 0.0001.

### Structural implication of NSP12 mutations on NSP12-NSP8-NSP7 RdRp complex formation

We further performed structural simulation modelling to examine whether the NSP12 P323L or P322L/G617S mutation impacted the NSP12-NSP8 interaction interface (Extended Data Fig. 3). This suggested that the P323L mutation of NSP12 enabled stabilized hydrophobic interactions with neighboring residues including F326, F396, and V675 (Extended Data Fig. 3a). This intramolecular interaction change may affect hydrophobic residues such as V115, P116, L117, and I119 of NSP8 inducing a stronger interfacial interaction between NSP12 and NSP8. Similarly, the G671S mutation of NSP12 may result in a local structural change of the loop, encompassing residues A400 to F407, via hydrogen bonding between the side chain hydroxyl group of G671S and the backbone amide nitrogen of A406 (Extended Data Fig. 3b). This predicted conformational change in the interface region may reinforce the interfacial interaction between NSP12 and NSP8, through strengthened hydrophobic interactions between residues L388, F407, and V405 of NSP12 and residues I185 and M129 of NSP8.

**Fig. 3.**
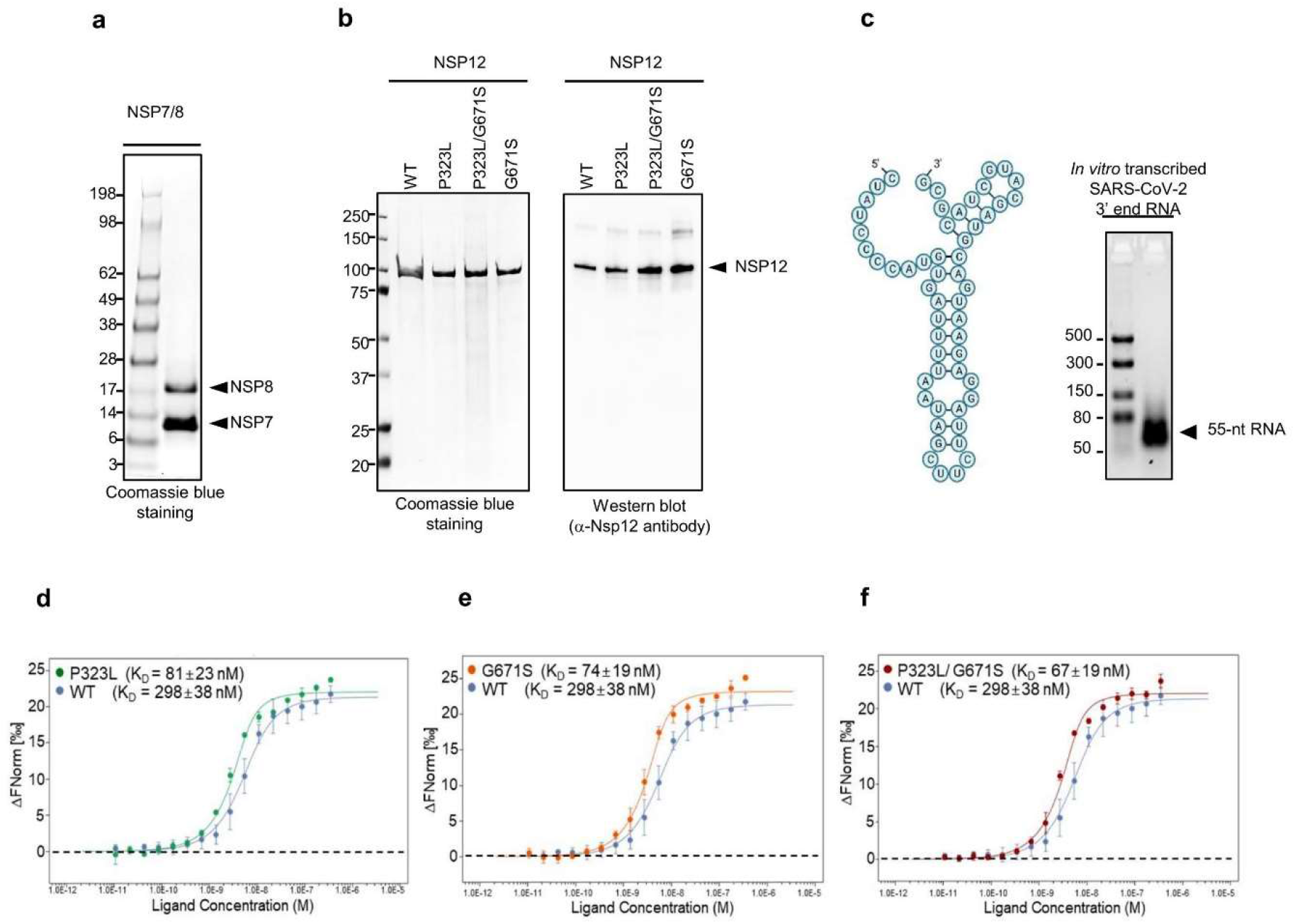
SARS-CoV-2 N12 bearing the P323L or P323L/G671S mutations enhances RdRp interaction with NSP7/NSP8/RNA. **a** SDS-PAGE of purified SARS-CoV-2 NSP7/NSP8confirmed by Coomassie blue staining. **b** SDS-PAGE of purified SARS-CoV-2 NSP12 WT, P323L, G671S, or P323L/G671S indicated by Coomassie blue staining (left) or western blot using anti-NSP12 antibody (right). **c** *In vitro* transcribed SARS-CoV-2 3′-end 55-nt RNA in 3% agarose gel (right). The predicted secondary structure of the SAR-CoV-2 3′-end 55-nt RNA (left). (**d, e,** and **f**) Microscale thermophoresis assay was performed to analyze the ability of RdRp to form a complex with experimental sets of **d** WT versus P323L, **e** WT versus P323L, **f** or WT versus P323L/G671S NSP12. Binding affinity was examined by monitoring the thermophoretic traces, and the dissociation constant (K_D_) was determined by measuring the change in fluorescence (ΔFnorm). Data are presented as the mean ± s.d. of three independent analyses.

Based on the structural analysis findings, we next used a highly sensitive microscale thermophoresis (MST) assay system ^22^ to test how the NSP12 mutations affected the stability of the NSP12-NSP8-NSP7 complex. Briefly, NSP7 and NSP8 proteins were purified from E. coli (Fig. 3a), and WT, P323L, and P323L/G671S NSP12 proteins were purified from Sf9 insect cells using baculovirus expression systems (Fig. 3b) ^8,23^. In vitro transcribed SARS-CoV-2 3′-end RNA ^23^ was used to quantify binding affinities of the NSP12-NSP7-NSP8-RNA complex (Fig. 3). We found that the dissociation constant (KD) to NSP7-NSP8-RNA was 298±38 nM for NSP12 WT, 81±23nM for P323L mutant, 74±19 nM for G671S mutant, and 67±19 nM for P323L/G671S mutant (Fig. 3d-f). These results indicate that the NSP12 P323L or P323L/G671S mutations play a pivotal role in promoting the stability of RdRp complex formation, leading to increased RdRp activity.

### The P323L and G671S mutations in NSP12 alter SARS-CoV-2 fitness and transmission

To evaluate how the P323L and G671S mutations affected SARS-CoV-2 fitness and transmission, we used the SARS-CoV-2 Bacmid reverse genetic (RG) system to generate three isogenic SARS-CoV-2 variants (Spike D614G) containing the NSP12 WT, P323L, G671S, or P323L/G617S mutations ^24^. We then examined their in vitro replication kinetics in Vero, Vero^ACE2/TMPRSS2^, Huh-7, and Calu-3 cells at 33°C or 37°C. Strikingly, in all four different cell types, RG-P323L and RG-P323L/G671S mutant viruses replicated at much higher viral titers than RG-WT virus, with more pronounced enhancement of viral titers at 33°C than at 37°C (Fig. 4). On the other hand, RG-G671S mutant virus showed replication similar to RG-WT virus, although it exhibited slightly higher virus titers than RG-WT virus in Huh-7 cells (Fig. 4e,f).

**Fig. 4.**
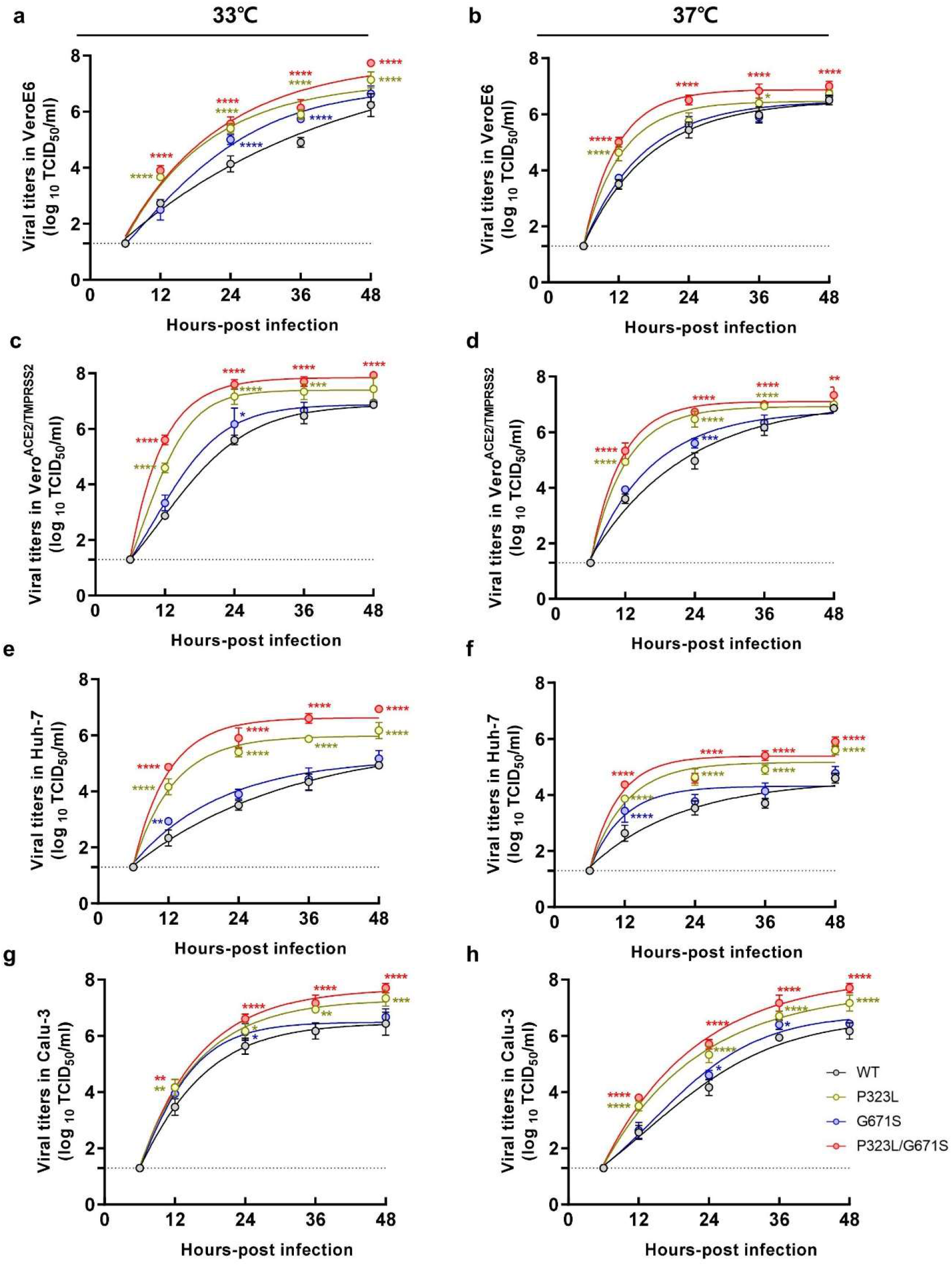
Growth kinetics of RG-derived SARS-CoV-2 *in vitro*. Comparison of differential growth kinetics among RG SARS-CoV-2 viruses (WT, P323L, G671S, and P323L/G671S mutant) in **a-b** Vero, **c-d** Vero^ACE2/TMPRSS2^, **e-f** Huh-7, and **g-h** Calu-3 cells at 33°C or 37°C. Cells were infected with each RG virus at an MOI of 0.01. The supernatants of infected cells were harvested at the indicated time-points, and virus titers were determined using tissue culture infection dose 50% (TCID_50_) in Vero cells. The *p* values were calculated using a two-way ANOVA, with Turkey’s post-test. \**p* < 0.05; \*\**p* < 0.01; \*\*\**p* < 0.001; \*\*\*\**p* < 0.0001.

To further evaluate how the NSP12 P323L and G671S mutations affected in vivo viral growth and transmission, 20- to 24-month-old ferrets were infected with equal titers of RG-WT, RG-P323L, RG-G671S, or RG-P323L/G671S mutant virus. In the direct infection groups, RG-P323L, RG-G671S, and RG-P323L/G671S mutant viruses consistently exhibited higher virus titers than RG-WT virus (Fig. 5a, left panel). Notably, compared to the other ferret groups, RG-P323L/G671S mutant-infected ferrets exhibited the highest viral titers in the nasal turbinates (Fig 5a). In the first-round transmission study, infectious viruses were recovered from the nasal wash specimens of all DC ferrets as early as 2 dpc. Compared to RG-WT virus, RG-P323L and RG-P323L/G671S viruses showed significantly higher titers, with longer virus shedding until 6 dpc. Notably, RG-P323L/G671S mutant virus consistently showed the highest virus titers (Fig. 5a). In the second-round transmission study, infectious RG-P323L/G671S virus was recovered in the nasal washes from all three second DC ferrets, as early as 2 dpc, with the highest viral titers found until 6 dpc. Compared to RG-WT virus, RG-P323L virus exhibited higher viral titers from 3 to 5 dpc (Fig. 5a, right panel). RG-G671S mutant virus showed attenuated virus titers compared to RG-P323L and RG-P323L/G671S mutant viruses but exhibited higher titers than RG-WT virus.

**Fig. 5.**
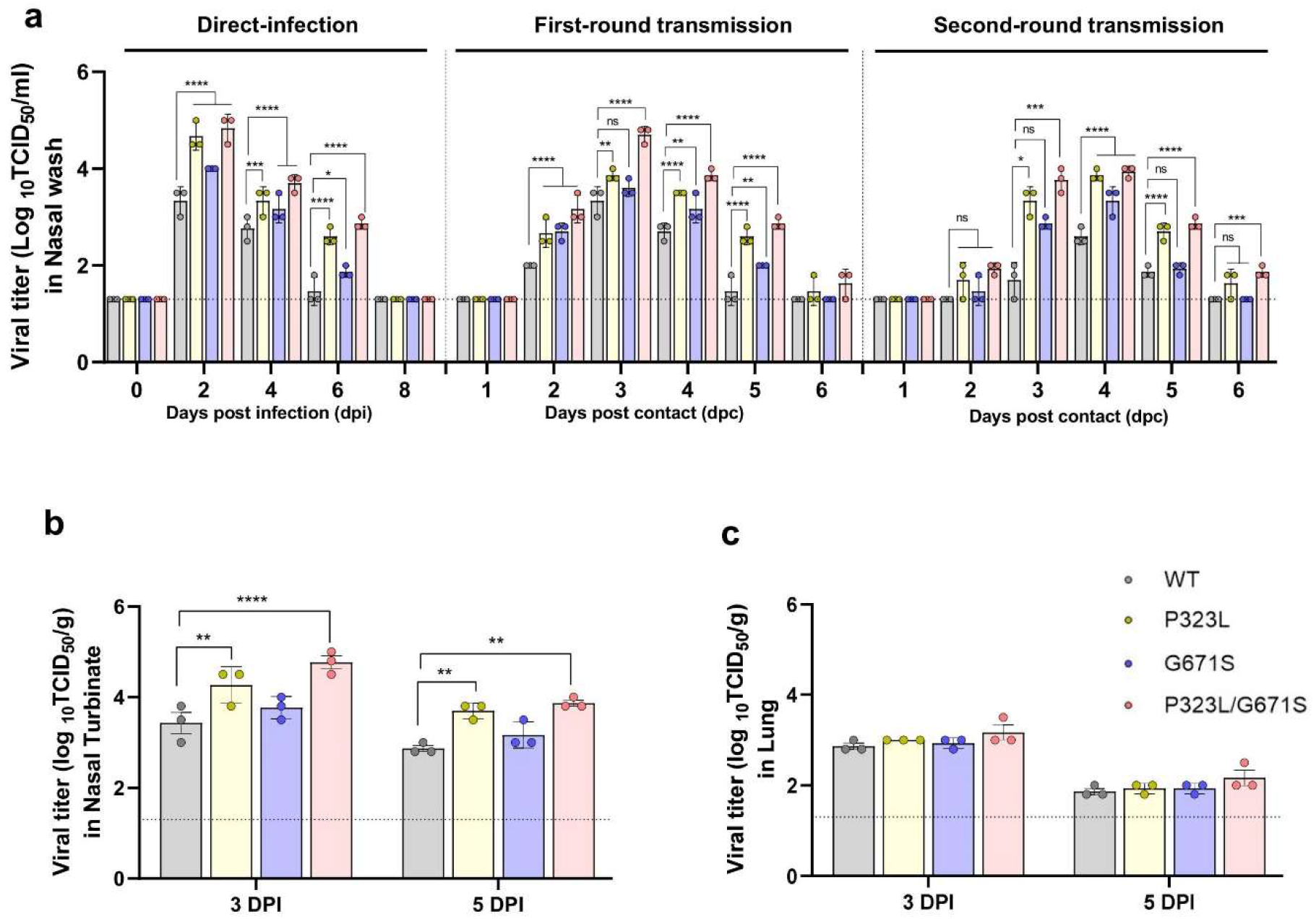
Replication and transmission of RG-derived SARS-CoV-2 in ferrets. Ferrets (*n* = 9/group) were inoculated intranasally with 10^5^ TCID_50_/ml of RG-derived SARS-CoV-2. Nasal wash specimens were collected at a 2-day interval from directly infected ferrets, and daily from first-round and second-round transmission ferrets. Ferrets (*n* = 3/group) were sacrificed at 3 and 5 dpi. The infectious virus titers in **a** nasal wash specimen, **b** nasal turbinates, and **c** lung tissue samples were measured by determining the TCID_50_ per unit of measurement (g or mL) using VeroE6 cells. The limit of detection (1.3 log_10_ TCID_50_/ml or log_10_ TCID_50_/g) is indicated by a dotted line for each representation. The *p* values were calculated using a two-way ANOVA, with Turkey’s post-test.\**p* < 0.05; \*\**p* < 0.01; \*\*\**p* < 0.001; \*\*\*\**p* < 0.0001.

To evaluate the in vivo replication properties of each mutant RG virus, directly infected ferrets (n = 3/group) were sacrificed at 3 and 5 dpi, and nasal turbinates and lung tissues were collected for viral titration. In the direct infection groups, compared to RG-WT virus, RG-P323L and RG-P323L/G671S mutant viruses consistently exhibited higher virus titers in the nasal turbinates at 3 and 5 dpi (Fig. 5b). In contrast, virus titers did not significantly differ in the lung tissues of all infected ferrets (Fig. 5c). Collectively, these results demonstrated that the P323L and P323L/G671S mutations of NSP12 enhanced virus growth properties in the upper respiratory tract, promoting rapid transmission to direct contact naïve animals.

## Discussion

The emergence and widespread transmission of SARS-CoV-2 variants has significantly impacted the COVID-19 pandemic, resulting in several waves and substantial surges of VOC infections worldwide. Epidemiological studies have reported that the rapid transmission of these VOCs is closely associated with mutations in the viral genome ^25–27^. However, systematic comparison studies are needed to compare the relative transmissibility of recent SARS-CoV-2 VOCs. In this study, we conducted systematic transmission studies in a ferret animal model, and our results revealed that the Alpha, Beta, and Delta variants exhibited rapid transmission to direct contact naïve ferrets, with high virus titers and prolonged viral shedding. Interestingly, only the virus titers in the nasal turbinates were significantly increased in ferrets infected with VOCs compared to the reference strain, while all strains led to comparable lung tissue virus titers. This suggests that natural temperature differences between the upper and lower respiratory tracts may have influenced the virus’ replication machinery.

Notably, recent SARS-CoV-2 VOCs commonly harbor the P323L mutation in NSP12, which encodes RdRp protein. RdRp plays a key role in viral RNA synthesis, together with NSP7 and NSP8, which function as co-factors in virus replication and transcription ^9^. Thus, we reasoned that the P323L mutation may lead to altered polymerase activity, which could be associated with high viral replication in the respiratory tracts. To better understand difference of infectious virus titers between nasal turbinates and lungs, we conducted RdRp reporter assay at 33°C or 37°C to mimic the upper and lower airway temperature, respectively. Notably, the NSP12-P323L mutation resulted in enhanced RdRp activity at 33°C but not at 37°C. Moreover, the highest polymerase activity was observed with the NSP12 P323L/G671S mutation found in Delta variants. These results suggest that the relatively low temperature (33°C), typical of the upper respiratory tract, promotes enhanced RdRp activity of the NSP12 P323L/G671S mutant, and thereby increases SARS-CoV-2 replication in the upper respiratory tract.

Structural simulation analysis of SARS-CoV-2 NSP12-NSP8 complex showed that NSP12 mutations may reinforce the interfacial interaction between NSP12 and NSP8 through strengthened hydrophobic interactions, resulting in improved assembly of the replication machinery for efficient viral replication and transmission. This hypothesis was substantiated by MST assay to verify enhanced NSP12-NSP8 complex interaction. This reveals that compared to NSP12 WT, NSP12 P323L or P323L-G671S mutant displays significantly increased interaction with NSP8, resulting in enhanced RdRp activity.

We also demonstrated that NSP12 mutations influenced in vivo replication kinetics in a temperature-dependent manner. Briefly, ferrets that were directly infected with the RG-P323L and RG-P323L/G671S mutant viruses exhibited significantly enhanced virus titers in nasal washes, which led to rapid transmission to naïve contact ferrets. However, these viruses showed similar titers to WT virus in the lung tissues, suggesting that the NSP12-P323L mutation is closely associated with high viral fitness in upper respiratory tract (33°C) rather than lower respiratory tract (37°C). Similarly, a mammalian host adaptation-associated RNA polymerase PB2 E627K mutation of avian influenza A virus directly alters polymerase enzymatic kinetics in a temperature-dependent manner ^28^. This suggests that NSP12 mutations in SARS-CoV-2 have evolved to have enhanced polymerase activity and high viral replication in the upper respiratory (33°C), which ultimately contributes the rapid spread of recent SARS-CoV-2 VOCs in human populations.

In this study, we demonstrate that the P323L or P323L/G671S mutation in the NSP12 results in enhanced transmissibility due to the increased viral growth properties and extended infectious virus shedding duration in the upper respiratory tract. Interestingly, all Omicron variants, from BA.1 to BA.5, commonly harbor the NSP12 P323L mutation ^29–31^ and BA.2.75 Omicron variants also carry the NSP12 P323L/G671S mutation ^32^, similar to the Delta variant. These NSP12 mutations may be responsible for high transmissibility of Omicron variants. Thus, our data unambiguously demonstrate that mutations in NSP12 increases viral replication in the upper respiratory tracts, which promotes the high transmissibility of SARS-CoV-2 VOC strains.

## Materials and Methods

### Basic sequence analysis and selection of SARS-CoV-2 viruses

SARS-CoV-2 viruses of each clade were isolated from clinical samples collected from COVID-19 patients at Chungbuk National University Hospital between February 2020 and March 2021. Each SARS-CoV-2 virus was isolated after three passages in Vero cells in DMEM (Sigma Aldrich). Virus isolation was confirmed by RT-PCR using E gene-specific primer sets (Extended Data Table 2). For next-generation sequencing (NGS), whole SARS-CoV-2 genome cDNA libraries were generated using a MiniSeq system (Illumina) with a MiniSeq mid-output kit, resulting in 2 × 150 nucleotides paired-end reads. For each SARS-CoV-2 clade, the complete genome sequence was determined in parallel to the reference genome Wuhan-Hu-1-like (also known as Clade L). Alignment of the sequences for each clade and recent variant genome was performed using the QIAGEN Bioinformatics CLC Workbench program (version 10.1.1; CLC bio). All described SARS-CoV-2 whole-genome sequences are available in GISAID (Extended Data Table 1).

### Cells and in vitro virus growth kinetics

Vero, Vero^ACE2/TMPRSS2^, Huh7, and Calu-3 cells were cultured in DMEM supplemented with 10% FBS and 1% penicillin-streptomycin. To establish Vero^ACE2/TMPRSS2^, Vero cells were infected with the lentiviral vectors pLL-CMV-hACE2-bla and pLL-CMV-hTMPRSS2-puro, using a Lenti-X Concentrator (Clontech Laboratories, Inc., Mountain View, CA, USA) following the manufacturer’s instructions. Each cell line was inoculated with each virus at 0.01 multiplicity of infection (MOI). Virus culture media was harvested at 12, 24, 36, and 48 hours post-infection (hpi), and were used to titrate Vero cells to determine the 50% tissue culture infective dose (TCID_50_). After 3 days, the cells were fixed with 10% formalin (v/v), and visualized by staining with 1% crystal violet (w/v).

### Generation of cell-based SARS-CoV-2 RdRp activity reporter assay

The cell-based reporter system included the bicistronic reporter construct containing firefly luciferase in the sense orientation, and Nano-glo® luciferase in the anti-sense orientation, which was flanked by the antisense 3′- and 5′-UTR of SARS-CoV-2 and the hepatitis D virus ribozyme (HDV) self-cleavage sequence. The reporter plasmid contained the sense firefly luciferase gene (GenBank®: U47295), hepatitis D virus ribozyme sequence, antisense 3′-untranslated region (UTR) of SARS-CoV-2, anti-sense Nano-glo® luciferase gene (NLuc) (GenBank®: KM359770), and HDV ribozyme sequence, which were synthesized in a row by GENEWIZ, and cloned into the HindIII and XhoI sites of the pcDNA3.1(+) plasmid (Invitrogen, Carlsbad, CA, USA). The C-terminal 10× His-tagged NSP12 gene, and N-terminal Flag-tagged NSP7 and NSP8 gene (GenBank®: NC_045512) were cloned into the BamHI and XhoI, and NheI and XhoI sites, respectively, of the pcDNA3.1(+) plasmid. For construction of NSP12 mutants (P323L, P323L/G617S, and G617S), site-directed mutagenesis was performed using the following primer sets (Extended Data Table 3). The 293T cell line was maintained in Dulbecco’s modified Eagle’s medium (Gibco) supplemented with 10% fetal bovine serum (FBS) and 1% penicillin-streptomycin, at 37°C in 5% CO_2_. For transient transfection, 293T cells were seeded in 96-well plates overnight. Plasmids were mixed at a ratio of 1:1:1, 1:2:1, or 2:2:1 (*33*) with TransIT®-LT1 transfection reagent (Mirus Bio LLC, Madison, WI, USA) and the plasmids. These reagent mixtures were added to the cells according to the manufacturer’s instructions, and incubated for 24 h at 33°C or 37°C in 5% CO_2_. In these cells, the Fluc and NLuc reporter gene expression was measured using a Nano-Glo® Dual-Luciferase® Reporter Assay System (Promega Corporation) following the manufacturer’s instructions.

### Purification of the recombinant SARS-CoV-2 NSP12 proteins

Recombinant SARS-CoV-2 NSP12 WT, P323L, G671S, and P323L/G671S N-terminally tagged with 6×His were purified using Bac-to-Bac™ Baculovirus Expression System based on the manufacturer’s instruction (Gibco). Upon 3 dpi of NSP12-enconding baculoviruses into Sf9 cells, the recombinant NSP12-containing culture supernatant from the baculovirus-infected Sf9 cells was pre-cleared by centrifugation. Then collected supernatant was loaded onto the HisPur™ Ni-NTA Spin Purification Kit (Thermo Scientific). Captured NSP12 proteins were eluted with an elution buffer containing 300 mM imidazole and the purified NSP12 proteins were dialyzed with a dialysis buffer containing 150 mM NaCl, 20 mM Tris (pH8.0), and 1 mM of 2-mercaptoethanol. Then, the purified proteins were aliquoted and stored at −80 °C until use.

### Purification of the recombinant SARS-CoV-2 NSP7 and NSP8 proteins

Recombinant NSP7 and NSP8 proteins were expressed and purified by a dual expression system using pCDFDuet vector as previously reported (23). Briefly, 500 ml LB culture of pCDFDuet-1-6×His-SARS-CoV-2-NSP7/NSP8 (Addgene)-transformed *E.coli* BL21 DE3 strain was induced by isopropyl β-d-1-thiogalactopyranoside (IPTG, 0.1 mM final) for 14 hr at 16 °C. Then, the cells were collected by centrifugation, resuspended in the buffer containing 20 mM Tris (pH 8.0), 0.1 mM EDTA (pH 8.0), 300 mM NaCl, 5% (v/v) glycerol, 5 mM imidazole, 1 mM 2-mercaptoethanol, 1 mM PMSF, and a protease inhibitor cocktail (Roche), lysed by sonication, and cleared by centrifugation. Finally, the recombinant NSP7/NSP8 was purified using a HisPur™ cobalt resin column (Cytiva). To remove N’ terminus 6×His at NSP7, a recombinant HRV 3C protease (Sigma-Aldrich) was added to NSP7/NSP8 complex and incubated overnight at 4 °C. Processed NSP7/NSP8 was purified by passing through the HisTrap HP column and the flow-through was collected and dialyzed overnight with a dialysis buffer containing 150 mM NaCl, 20 mM Tris (pH8.0), and 1 mM of 2-mercaptoethanol. Finally, purified proteins were aliquoted and stored at −80 °C until use.

### Generation of the SARS-CoV-2 3’ end 55-mer RNA by in vitro transcription

SARS-CoV-2 3’ end RNA was synthesized by in vitro transcription using an mMESSAGE mMACHINE T7 kit (Invitrogen) with a DNA template (5’- TAATACGACTCACTATAGCTATCCCCATGTGATTTTAATAGCTTCTTAGGAGAAUGACGTAGCATGCTACGCG -3’) according to the manufacturer’s instructions. The T7 promoter sequence in the DNA template is underlined.

### Microscale thermophoresis assay

Purified WT or mutant NSP12 recombinant proteins were labeled with Cytidine-5 dye using the Monolith His-Tag labeling kit (NanoTemper Technologies), and then labeled NSP12 proteins were diluted to 700 nM in PBST buffer. In parallel, the NSP7 (700 nM)-NSP8 (1.4 μM)-RNA (700 nM) mixture was prepared and diluted to make 16 tubes of 2-fold serial diluents with PBST buffer. For the binding experiment, 10 μl of the labeled NSP12 was mixed with 10 μl of each serial diluent. Then, the mixture was incubated at room temperature for 20 min, followed by centrifugation at 10,000 × g for 10 min. The mixed samples were then loaded into the Monolith NT.115 capillaries (NanoTemper Technologies) and the measurements were performed using a Monolith Pico instrument (NanoTemper Technologies) under the optimized experimental condition (50% excitation power, medium MST power, and ambient temperature of 25 °C). For the analysis of the RdRp complexability, the thermophoretic traces were monitored and the dissociation constant (K_D_) was determined by measuring the change in fluorescence (ΔFnorm). The results were plotted and the binding affinities were calculated by MO. Affinity Analysis software (NanoTemper Technologies).

### Animal study design

For ferret-to-ferret transmission, 20- to 24-month-old female ferrets (*n* = 9/group) were inoculated with 5.0 log_10_ TCID_50_/mL via the intranasal route, using each of the tested SARS-CoV-2 representative strains and variants, and the RG viruses. For the first-contact transmission study, at one day post-infection, one directly infected ferret from each group was randomly selected to be co-housed in direct contact with naïve ferrets (*n* = 3/group), as the first-round direct contact. For the second-round direct transmission study, at two days post-contact, the three first-contact ferrets were co-housed in direct contact with new naïve ferrets (*n* = 3/group). Following inoculation of the infected ferret groups, nasal wash specimens were collected every two days for eight days to measure virus shedding. To assess virus replication in different organs, directly infected ferrets (*n* = 3/group) were sacrificed at 3 and 5 dpi, and nasal turbinates and lung tissues were collected using individual scissors to prevent cross-contamination. From the first- and second-round direct transmission ferrets, nasal wash specimens were collected every day until 7 dpc. Infectious virus titers were measured in the nasal washes and homogenized tissues by determining the numbers of TCID_50_ per unit of measurement (g or mL) using VeroE6 cells.

### Ethics statement

All animal experiments were approved by the Medical Research Institute, a member of the Laboratory Animal Research Center of Chungbuk National University (LARC) (approval number CBNUA-1731-22-01) and were conducted in strict accordance with and adherence to relevant policies regarding animal handling as mandated under the Guidelines for Animal Use and Care of the Korea Centers for Disease Control (KCDC). The handling of virus was performed in an enhanced biosafety level 3 (BSL3) containment laboratory as approved by the Korea Centers for Disease Control and Prevention (protocol KCDC-14-3-07).

## Supporting information

supplementary data contained 3 figures and 3 tables.

## Acknowledgements

This work was supported by the Institute for Basic Science (IBS-R801-D1) to YKC; NIH CA200422, CA251275, AI140718, AI171443, AI171201, DE023926, DE028521 to JUJ; and AI140705, AI140705S, AI152190 to YKC and JUJ; and the National Research Council of Science & Technology (NST) grant by the Korea government (MSIT) (No. CAP20013-000) to MHK and EHK.

## Author contributions

S.M. Kim, E.H. Kim, J.U. Jung, and Y.K. Choi conceptualized and designed the study. E.H. Kim, S.M. Kim, Y.I. Kim, R. Sun, M. Kwak, J.S. Yoo, M.A. Yu, K.M. Yu, M.A.B. Casel, R. Rollon, S.G. Jang, J.H. Choi, J.R. Gil, K.Y. Eun, H.G. Kim, A. Ensser, J. Hwang, M.H. Kim, and Y.K. Choi undertook the investigation, and the data and viral genome sequencing analysis and interpretation. S.M. Kim, E.H. Kim, M.A.B. Casel, J.U. Jung, and Y.K. Choi drafted and collectively finalized the manuscript.

