## supplementary data contained 3 figures and 3 tables. for "SARS-CoV-2 variants show temperature-dependent enhanced polymerase activity in the upper respiratory tract and high transmissibility"

**Extended Data Fig. 1.**

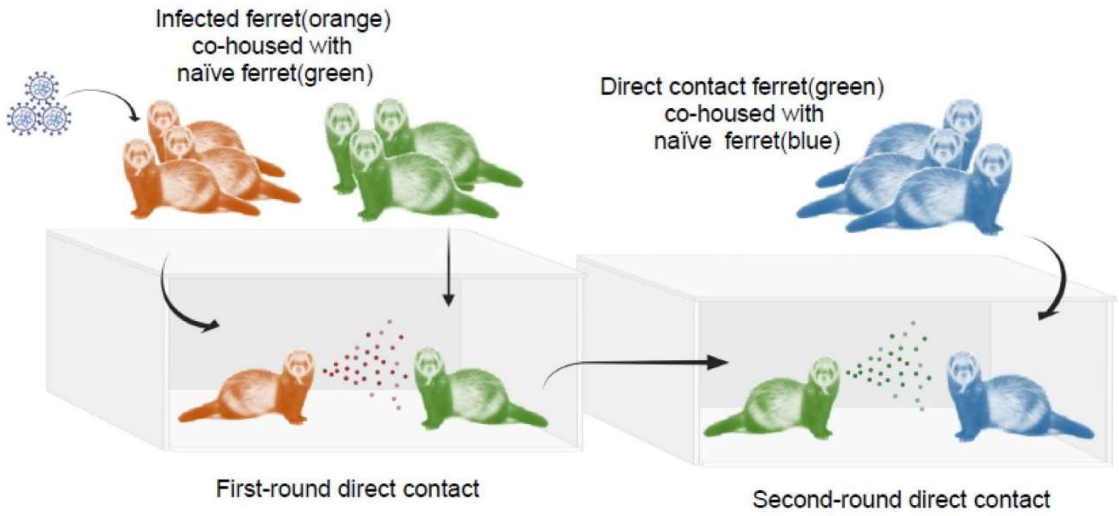


**Extended Data Fig. 1**. **Study design for animal-to-animal transmission in ferrets.**

For the first-round direct contact transmission study, one randomly selected directly infected ferret was co-housed with naïve contact ferrets (*n* = 3/group) at one day post-infection, and direct transmission was monitored. For the second-round direct contact transmission study, the first-round direct contact ferrets (*n* = 3) were introduced to new naïve ferrets (*n* = 3/group) two days after having been co-housed with directly infected ferrets. From the second-round direct contact ferrets, nasal wash specimens were collected every day for 7 days from each group.

Extended Data Fig. 2.

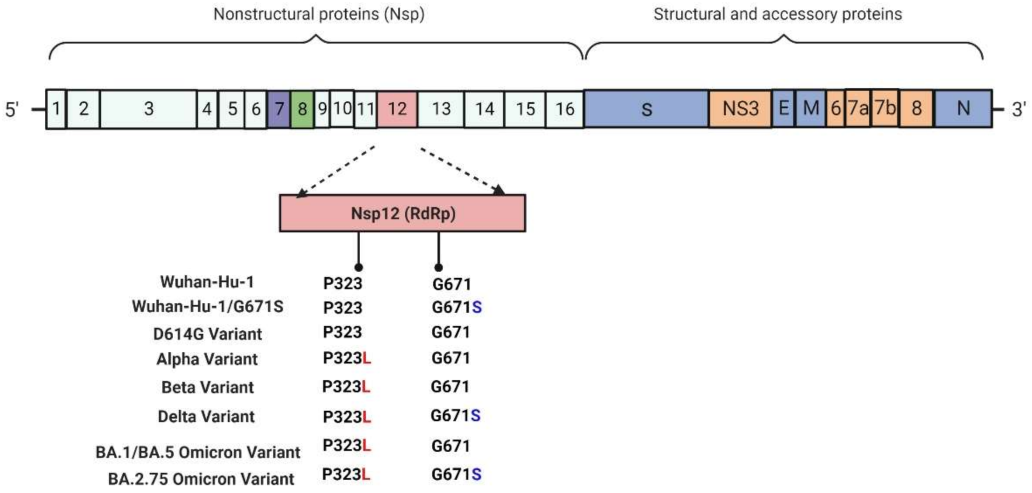


**Extended Data Fig. 2**. **SARS-CoV-2 NSP12 non-synonymous mutations.**

The results of amino acid genetic variance analysis summarized as a graph, showing information regarding genetic variations in the NSP12 among D614G and SARS-CoV-2 variants in comparison with the Wuhan-hu-1 (Clade L) reference sequence.

Extended Data Fig. 3.

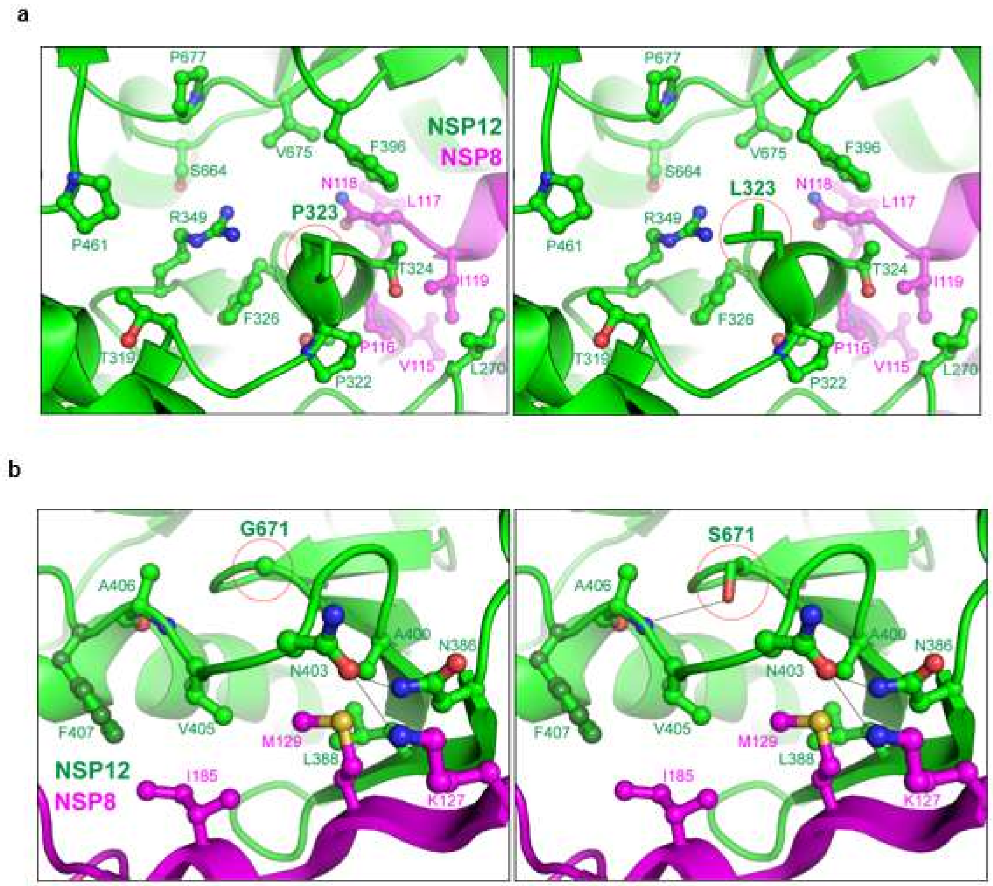


**Extended Data Fig. 3**. **Structural analysis of the effects of the P323L and G671S variations on the NSP12 and NSP8 complex.**

**a**. Pro323 (red dashed circle) and neighboring residues in the NSP12 replication and transcription complex (RTC) structure (PDB entry 7CYQ, left panel). The mutation of Pro323 to leucine is shown in the red dashed circle. The estimated interaction residues of NSP12 and NSP8 are illustrated with green and magenta, respectively (right panel). **b**. Gly671 (red dashed circle, left panel) and the mutation of Gly671 to serine (red dashed circle, right panel) are shown in the same SARS-CoV-2 RTC structure, and the neighboring residues are illustrated. The structure models showing the mutations were generated using PyMOL (Schrödinger, L. & DeLano, W., 2020. *PyMOL*, Available at: http://www.pymol.org/pymol.). NSP12 and NSP8 are represented in green and magenta, respectively. Hydrogen bonds are indicated as black dashed lines.

Extended Data Table 1. **Sequence variants of SARS-CoV-2 clade named by GISAID.**

| **GISAID Clade** | **PANGOLIN lineage** | **Substitutions** | **Remark** |
|---|---|---|---|
| L | - | - | Reference |
| G | B.1 (D614G) | ORF1ab: C241T, C3037T; Spike: D614G | CBNU-nCoV 05 |
| GRY | B.1.1.7 (Alpha) | ORF1ab: C241T, C3037T; NSP3: T183I, A890D, S1038F, I1412T; NSP5: K90R; NSP6: S106K, Δ107–108; NSP12: P323L; Spike: Δ69–70, Δ144, N501Y, A570D, D614G, P681H, T716I, S982A, D1118H; ORF9: Q52I, Y73C; N: D3L, R203K, G204R, S235F | CBNU-nCoV 11 (UK variant-like) |
| GH/ 501Y.V2 | B.1.351 (Beta) | ORF1ab: C241T, C3037T; NSP3: V473F, L620F, S794L; NSP5: K90R; NSP6: S106K, Δ107–109; NSP12: P323L; Spike: L18F, L54F, D80A, D215G, Δ242–244, K417N, E484K, N501Y, D614G, A701V; E: P71L, N:T205I | CBNU-nCoV 21 (South African variant-like) |
| GK G/452R.V3 | B.1.617.2 (Delta) | ORF1ab: C241T, C3037T; NSP3: A489S, A890D, P1228L; NSP12: P323L, G671S; NSP13: P78L; Spike: T19R, G142D, E154K, L452R, E484K, D614G, P681R, D950N, Q1071H; ORF8: D119I, Δ120–121; N: D63G, R203M, D377Y | CBNU-nCoV 36 (India variant-like) |

Extended Data Table 2. **Primer sets used to confirm virus isolation and qRT-PCR.**

| | **Target gene** | **Forward Primers (5′→3′)** | **Reverse Primers (5′→3′)** |
|---|---|---|---|
| Virus isolation | E | ATGTACTCATTCGTT TCGGAAGAG | TTAGACCAGAAGATCAGG AACTCTAG |
| qRT-PCR | S | AGGGCAAACTGGAAAGA TTGCTGA | TCTGTGCAGTTAACATCCT GATAAAGAAC |

Extended Data Table 3. **Primer sets used to generate SARS-CoV-2 NSP12, NSP7, and NSP8 genes.**

| **Gene** | | **Forward Primers (5′→3′)** | **Reverse Primers (5′→3′)** |
|---|---|---|---|
| NSP12 | P323 | GCTAGCTCAGCTGATGCACAATCG | CCGCTCGAGCATCATCATCATCATCATCATCATCATCATCTGTAAGACTGTATG |
| | P323L | CTC TAC AGT GTT CCC ACT TAC AAG T | AAGTGGGAACACTGTAGAGAATAAAACATT |
| | G671S | TCA TGT GTG GCG GTT CAC TAT ATG T | ACC GCC ACA CAT GAC CAT TTC |
| NSP7 | | CCGGCTAGCGACTACAAAGACGATGACGACAAGTCTAAAATGTCAGAT | CCGCTCGAGTTGTAAGGTTGCC |
| NSP8 | | CCGGCTAGCGACTACAAAGACGATGACGACAAGGCTATAGCCTCAGAG | CCGCTCGAGCTGTAATTTGACAGC |
